# Drivers of an epic radiation: the role of climate and islands in species diversification and reproductive-mode evolution of Old-World tree frogs

**DOI:** 10.1101/2021.09.22.461377

**Authors:** Gajaba Ellepola, Marcio R. Pie, Rohan Pethiyagoda, James Hanken, Madhava Meegaskumbura

**Author notes:** Corresponding author (MM).

## Abstract

Although large diversifications of species occur unevenly across space and evolutionary lineages, the relative importance of their driving mechanisms, such as climate, ecological opportunity and key innovations, remains poorly understood. Here, we explore the remarkable diversification of rhacophorid frogs, which represent six percent of global amphibian diversity, utilize four distinct reproductive modes, and span a climatically variable area across mainland Asia, associated continental islands, and Africa. Using a complete species-level phylogeny, we find near-constant diversification rates but a highly uneven distribution of species richness. Montane regions on islands and some mainland regions have higher phylogenetic diversity and unique assemblages of taxa; we identify these as cool-wet refugia. Starting from a centre of origin, rhacophorids reached these distant refugia by adapting to new climatic conditions (‘niche evolution’-dominant), especially following the origin of key innovations such as terrestrial reproduction (in the Late Eocene) or by dispersal during periods of favourable climate (‘niche conservatism’-dominant).

## Introduction

Since Darwin, understanding the processes underlying large-scale diversifications, wherein large assemblages of closely related lineages evolve from a common ancestor, have inspired evolutionary analyses^1–4^. Although several factors may mediate diversification, including geography, ecological opportunity, and key innovations^5^, the role of the ecological niche, particularly its climatic axes, remains poorly understood^6–9^.

Two main processes are central to understanding the role of climatic niche in the evolution of geographical distributions: niche conservatism (NC), where species tend to maintain their ancestral climatic niche over time^10-13^; and niche evolution (NE), where species adapt to new climatic conditions^7-9^. To colonize climatically similar areas when NC predominates, a species requires optimal climatic conditions to move across erstwhile climatic barriers^14^, whereas if NE predominates, species adapting to new conditions can overcome such barriers. Hence, knowledge of how climatic niches change over evolutionary timescales can enhance our understanding of the current distribution of lineages^8^.

Rhacophorid tree frogs comprise a spectacular diversification that can be used to test climatic correlates of evolution. Encompassing nearly 6% of global anuran (frog and toad) diversity, with 432 species in 22 genera^15,16^, rhacophorids occupy a large and climatically variable geographic area, mainly in Asia but with a single clade in Africa^15-18^. Across this range, rhacophorids are dispersed in distinct biogeographic regions^17,18^, which include swaths of continental mainland (East/Southeast Asia, peninsular India and Africa), continental islands (Japan, Taiwan, Hong Kong, Hainan, Sri Lanka and Andaman Islands), archipelagos (Sundaland, the Philippines), and montane regions (Himalaya). The northeastern region of the subtropical-temperate Asian mainland and islands holds the early-emerging genera, which are thought to have evolved 68–53 mya^17^.

Rhacophorid diversity is clustered spatially and temporally^17–20^. Some regions have greater clade or genus-level diversity (i.e., higher-level diversity), while others have high species diversity. Yet others are depauperate in both. Areas of high species diversity are often regarded as sources of diversification (species pumps), which nourish adjacent areas with lineages that evolved *in situ*^21^. However, regions with high diversification rates^22^ are not always those with high phylogenetic diversity, because species diversification may occur within just one or only a few clades or genera. Although highly diverse regions exhibit distinctive as well as shared climatic characteristics, the processes that mediate rhacophorid diversification, and especially the role of climate, remain poorly known.

Understanding of this is made more complicated, however, by rhacophorids being reproductively diverse, exhibiting four distinct modes: aquatic breeding (AQ; gel-covered aquatic eggs and aquatic tadpoles), gel nesting (GN; gel-covered terrestrial eggs and aquatic tadpoles), foam nesting (FN; foam-covered terrestrial eggs and aquatic tadpoles) and direct development (DD; terrestrial eggs and froglets fully developed at hatching)^18,19,23^. The likely ancestral mode of nearly all currently recognized genera (except the basal, fully aquatic forms) is gel nesting, while DD and FN appear to be key innovations that enabled rapid diversification^19^. Because geographic ranges of some genera exhibiting the latter two reproductive modes (FN, DD) do not overlap, each mode may represent an evolutionary response to variable climatic conditions in the past. However, the climatic correlates of these reproductive modes are poorly understood and some of them may represent adaptations to spread across climatic barriers (in the context of NE).

Diversification of ectothermic tetrapods, such as anurans, is influenced by climate^24^. Since rhacophorids are distributed across a climatically variable range, and given that their diversity is clustered in space and time^17,18^, we hypothesize that their diversification and dispersal has a strong climatic context (NE vs. NC), which is also associated with reproductive mode evolution. This hypothesis generates several predictions: (1) Rhacophorid diversity is clustered in climatically similar regions. A clade-level analysis should show the extent to which derived clades have shifted from the climatic niches of early-emerging clades as well as the role of NE vs. NC for each clade. (2) Regions with relatively high diversification rates such as islands and archipelagos have increased ecological opportunity, and are characterized by reproductive modes regarded as key innovations (DD and FN). (3) NE is achieved through reproductive mode evolution if correlated with specific climatic events. Analysis of this prediction will identify reproductive modes that enabled dispersal and diversification into optimal climatic oases such as islands and regions that are geographically disjunct from the centre of origin.

## Results and discussion

We constructed an updated phylogeny of the Rhacophoridae by including 415 extant species representing all 22 valid genera (Fig. 1). This represents the most complete taxon sampling of rhacophorids used to date, which enhances the accuracy and support for hypotheses testing of evolutionary relationships^25^. Our tree is congruent with many recent clade-level analyses (Supplementary Table 1); it offers relatively high support for most major nodes. It also resolves several long-standing taxonomic discordances within Rhacophoridae^18^. *Buergeria, Liuixalus, Theloderma* and *Nyctixalus* constitute early diverging lineages, while the remaining genera diverge into two major clades: clade A—*Beddomixalus, Gracixalus, Kurixalus, Mercurana, Nasutixalus, Philautus, Pseudophilautus* and *Raorchestes*; and clade B—*Chirixalus, Chiromantis, Feihyla, Ghatixalus, Leptomantis, Polypedates, Rhacophorus, Rohanixalus, Taruga* and *Zhangixalus*. Clade A is distributed largely across East and Southeast Asia, Sundaland and the Indian subcontinent, whereas clade B is distributed largely across East and Southeast Asia, Sundaland and Africa (Fig. 1).

**Figure 1.**
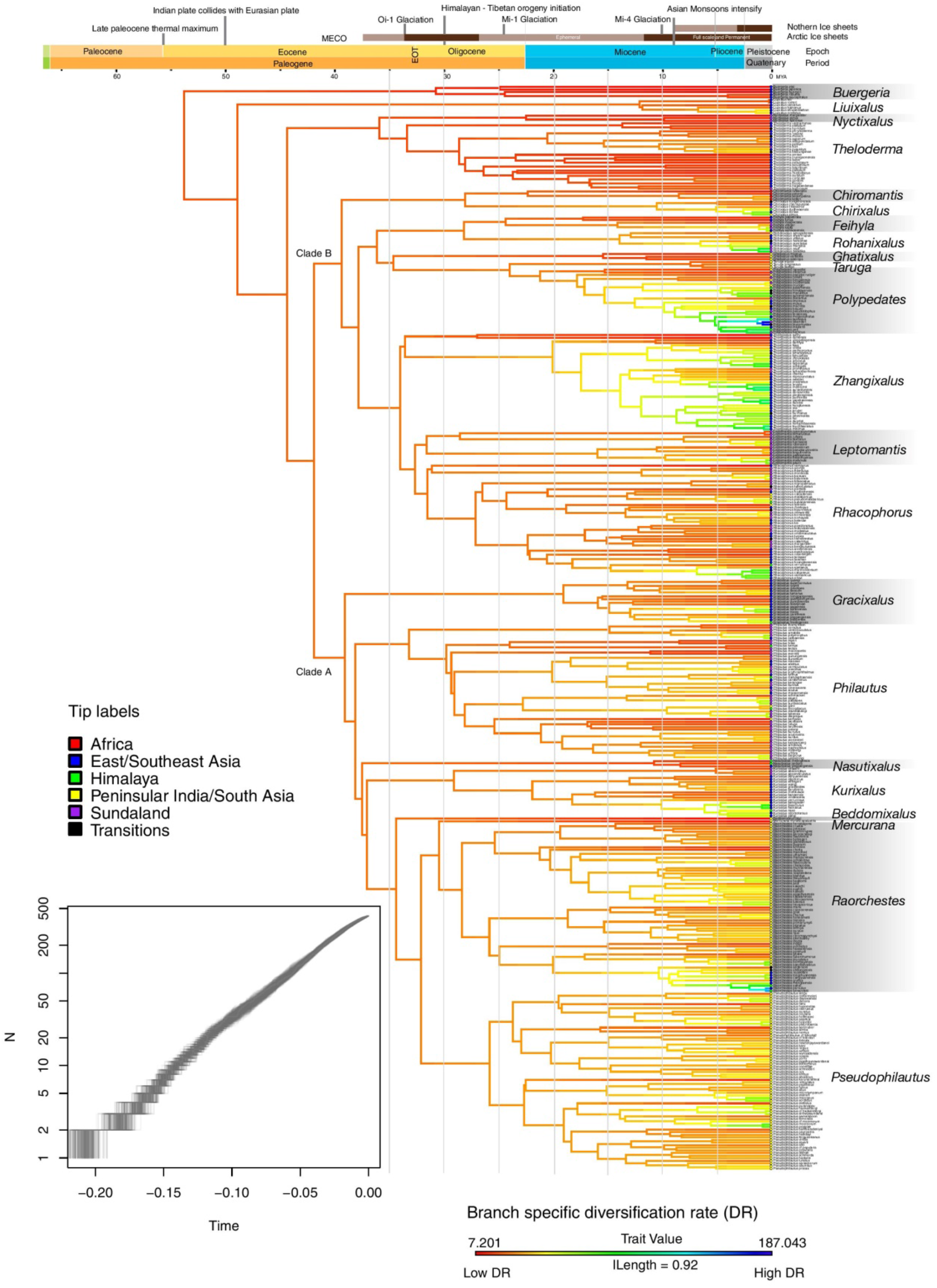
Complete phylogeny of rhacophorid frogs, with diversification rates (DR) traced across branches. The phylogeny is based on 58 AHE^18^ and 315 Sanger sequence data for 415 species. Molecular data are unavailable for 94 species, which are constrained by assessing other taxonomic information. The lineage-through-time plot (LTT; lower left) shows a constant rate of diversification but rates traced on the phylogeny show localized variation, especially among clades (genera): younger taxa have faster diversification rates (cool colours) than older, basal taxa (warmer colours). Colored dots at branch tips indicate the geographic region in which each species occurs; “Transitions” denotes species occurring in two or more regions.

Rhacophorid diversity is clustered spatially and temporally. Islands and some mainland regions have higher diversity and unique assemblages of taxa (Fig. 2a). In particular, species richness (SR) and phylogenetic diversity (PD) are highest in (1) montane and lowland rainforest areas of Borneo; (2) rainforest areas of peninsular Malaysia; (3) rainforest areas in the southern and northern Annamites of Vietnam; (4) northern Indochinese subtropical forests around Yunnan; (5) northern and southern montane rainforests of the Western Ghats in India; and (6) montane and lowland rainforests of Sri Lanka. These regions record more than 15 species per 1°×1° global grid cell (Fig. 2a). High correlations between SR and PD within these regions highlights their importance as species pumps and refugia for the family as a whole (Supplementary Fig. 1). Indeed, these regions are widely recognized as global centres of biodiversity^26–28^, especially amphibian diversity^29,30^.

**Figure 2.**
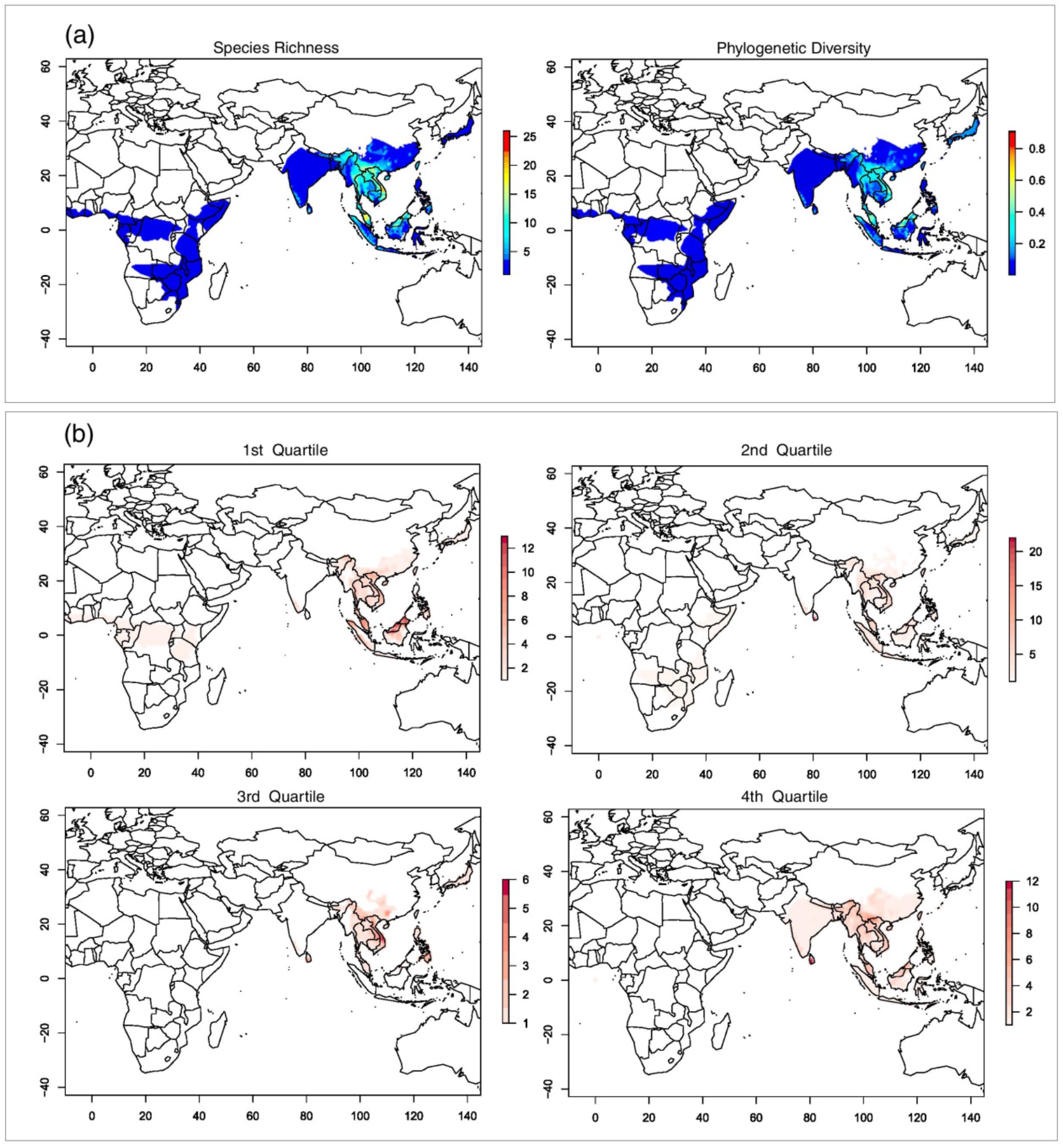
Patterns of spatiotemporal distribution in Rhacophoridae. (a) Spatial clustering of species richness (SR) and phylogenetic diversity (PD) in 1°×1° grid cells based on presence-absence matrices derived from species’ geographic ranges (see Methods). SR and PD are highest in the Sunda islands, peninsular Malaysia, Vietnam, the Yunnan-Guizhou plateau area, the Western Ghats of India and Sri Lanka. (b) Species richness in 1°×1° grid cells for each DR quartile. Dark red highlights grid cells with high species richness; light red indicates cells with low species richness within the respective DR quartile. DR values represent the number and timing of diversification events along a lineage: the 1^st^ DR quartile represents species with low diversification rates (consisting of older lineages having few close relatives), whereas the 4^th^ quartile represents species with faster diversification (having many young and close relatives). Highest species richness is observed during the 2^nd^ quartile and generally in the same regions as those of 1^st^-quartile species, indicating early dispersal and diversification events in confined areas in East/Southeast Asia, peninsular India and the Sundaland region. Regions of high SR and PD contain both the oldest and the youngest lineages. These regions may have served as species pumps and refuges during rhacophorid diversification.

The effect of climate on diversification is mediated by other factors such as the range of ecological opportunities and the particular lineages present at a given location, and thus has a strong geographic context^31^. However, geography alone cannot discern the role of climate, as similar climatic conditions may occur at different locations at different time periods and affect the evolution of independent lineages. Therefore, to understand patterns of occupation of climatic niche space and understand the climatic conditions under which rhacophorid species evolved, we extracted the mean climatic conditions of 3846 georeferenced coordinates of collecting localities of all species using data for 19 bioclimatic variables from WORLDCLIM 2.0^32^. We then used the average bioclimatic conditions in a principal component analysis (PCA) based on their correlation matrix, assuming that calculated species means reasonably approximate the realized climatic niche of a given species^9,33^.

The most important dimension of the rhacophorid climatic niche (Supplementary Table 2) is dominated by variation in temperature (PC1), particularly during the cold season (BIO-6), followed by mean temperature of the warmest quarter (BIO-10) and precipitation seasonality (BIO-15). Species from regions with the greatest SR and PD primarily occupy a realized climatic niche with cool-wet conditions (Fig. 3). Such conditions, marked by mild temperatures and humid climates, are prevalent in mainland (e.g., subtropical China) and mountainous regions in tropical islands characterized by monsoonal climates^34^. In short, rhacophorid diversity is clustered in climatically similar but geographically dissimilar regions (Figs. 2a, 3). As these areas are characterized by high PD^35^ as well, they also may have acted as refugia during rhacophorid diversification.

**Figure 3.**
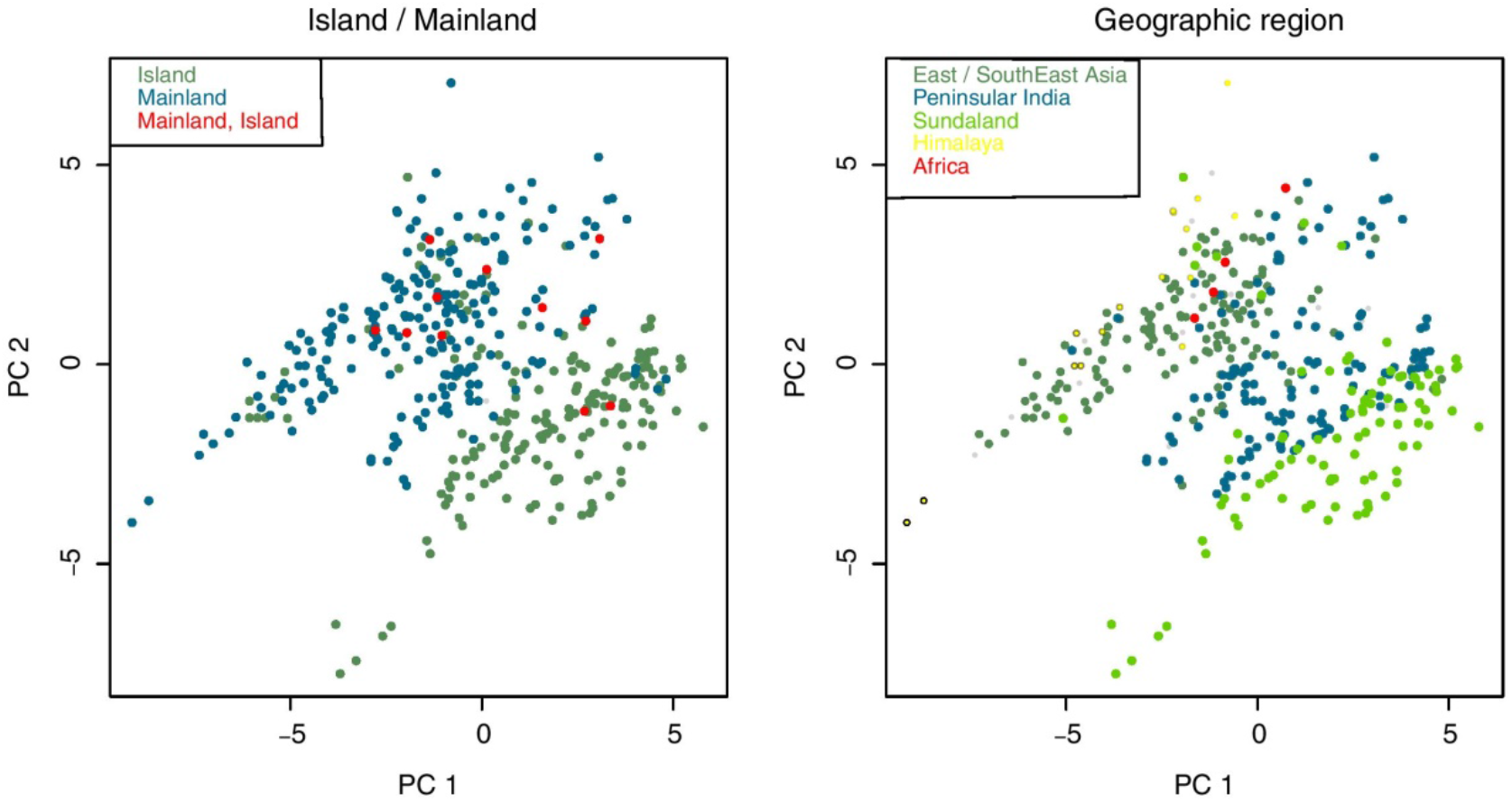
Climatic space defined by the first two principal component axes of climate niches in Rhacophoridae. Each point represents the average climatic conditions of a single species. Loadings are provided in Table S2. PC1 explains 40.5% of the variance and reflects variation in temperature; PC2 explains 23.9% of the variance and reflects variation in temperature and rainfall. Geographically, species in Sundaland are adapted to warm and less-seasonal climates, whereas species from East/Southeast Asia and Himalaya are cool-wet adapted. General patterns indicate that island species are comparatively warm adapted while mainland species are more or less cool-wet adapted.

Diversification rates (DR) overlain on the phylogeny show localized variation, especially in relation to clades: rates are low in phylogenetically isolated lineages resulting from early speciation events (i.e., basal lineages), whereas high rates are associated with members of terminal diverse lineages that originated from more recent speciation events (Fig. 1). We scored and ranked all lineages with respect to their relative phylogenetic isolation^3^ and then, based on these rankings, divided DR into quartiles in which the 1^st^ quartile contains the oldest and least diverse lineages and the 4^th^ quartile contains the youngest and most diverse ones (Supplementary Fig. 2). Subsequent mapping of species richness at a scale of 1°×1° for each quartile using IUCN species distribution range polygons^36,37^ reveals geographic variation in lineage accumulation through space and time (Fig. 2b). Furthermore, as the metrics SR, PD, DR and species crown age (SA) are significantly correlated (Supplementary Fig. 1), the spatial analyses yield similar patterns regardless of which metric is used. Both early-diverging and more recent lineages accumulate within the same regions, in which SR and PD are high. Based on ancestral geographic range reconstructions, Li et al. (2013)^17^ and Chen et al. (2020)^18^ concluded that mainland Asia played a significant role in the early diversification of rhacophorids and that ancestors of all early lineages in the Himalaya, peninsular India, Africa and Sundaland arrived via dispersal from mainland Asia, mostly in the Oligocene. Their conclusion is supported by our finding that species richness of older lineages is highest in East/Southeast Asia. We further suggest, given the high PD in these regions, that they might also have acted as refuges for Rhacophoridae, especially during early phases of its evolutionary history.

Reconstruction of variation in speciation rate (DR statistic) onto the maximum clade credibility tree shows that rates have been approximately constant throughout rhacophorid history, with only a few instances of rate increase or decrease towards the Miocene. This pattern is confirmed by the LTT plot based on 1000 post-burnin trees (Fig. 1). If climatic conditions within East/Southeast Asia were favourable during the period of origin of the Rhacophoridae, then one might instead expect to see an early burst of diversification^18,38^. We therefore suggest that early climates were less favourable for diversification of these frogs. This is supported by the fact that the common ancestor of the Asian tree frogs likely arrived from Madagascar around 60–70 mya possibly via the Indian plate^17^ (geological evidence for early India-Asia contact is still lacking), during a relatively warmer, climatically less favourable period^39^. However, assuming that the climatic conditions in which a species occurs reflects the conditions under which it evolved (viz., if NC predominated during early stages), rhacophorids appear to have survived in cool-wet, humid subtropical climates^40^, mostly towards the northeastern periphery of the family’s contemporary distribution. Moreover, as most early lineages occur in close proximity to their regions of origin (cool-wet climates), we infer that NC played a major role in early diversification (Fig. 2b; Q1). This claim is corroborated by the fact that the most recent common ancestor (MRCA) and older lineages of Rhacophoridae were aquatic breeders (AQ) ^18,19^, a life history mode which, in rhacophorids, depends on humid and cool climatic conditions.

Ecological opportunity would have been a significant factor enabling early dispersion and diversification of rhacophorids following their arrival on the Asian plate. Indeed, they dispersed extensively from their region of origin to other parts of Asia, India, Africa and Himalaya, as well as subtropical and tropical continental islands such as Taiwan, Japan, Sundaland, Philippines and Sri Lanka^18^ during phases of lowered sea level. Areas with low species richness of old lineages in the 1^st^ quartile (Fig. 2b) may have experienced extinctions and/or immigration of lineages that had already diversified elsewhere. The distinct increase in diversifications within the same regions of origin in the 2^nd^ quartile reaffirms the idea that NC was dominant during early dispersal and diversification.

Distribution-range expansion suggests that rhacophorids dispersed widely from their region of origin (North/Northeast Asia marked by cool-wet climates^40^) to adjacent regions during favourable climatic conditions for dispersal in the second and fourth quartiles of their diversification (Fig. 2b). Such climates have prevailed mainly during the Eocene-Oligocene transition (23–33 mya) and late Miocene (5.3–11.6 mya), during which time periodic glaciation led to depressed sea levels and the emergence of land bridges^39,41^, or when lowland dry zones became colder, enabling long distance dispersal of rhacophorids, followed by subsequent diversification^38,42^. Dispersals associated with niche conservatism might have been further facilitated by increased rainfall associated with the staged, rapid rise of the Himalaya and consequent strengthening of the Asian monsoonal cycle^20,43-45^.

Dispersal of rhacophorids from their centre of origin (cool-wet) to mainland (cooler, seasonal climates) and island refuges (warmer, less-seasonal climates; Fig. 3) requires traversing climatically harsh regions^46^ (especially for dispersal to peninsular India and Africa) and/or sea passages that appear intermittently via land bridges, as in the Sundaland region and between India and Sri Lanka. These intervening areas and land bridges are usually lowland areas with warmer climates. Therefore, apart from dispersing only during favourable periods (NC), crossing such areas periodically would also require adapting to harsh climates (NE). Indeed, the groups most successful at dispersal and habitat utilization did so in association with evolution of their climatic niche. Climatic correlates of evolution using ES-sim are not statistically significant (Supplementary Table 3), but other analyses suggest that climate played a major role in species diversification. For example, higher disparities shown along PC3 (dominated by high summer temperatures) during the 1^st^ quartile of diversification in East/Southeast Asia provide initial evidence that after the MRCA colonized this region, its descendants adapted (NE) to increasing summer temperatures (Fig. 4). The corresponding time periods coincide with the Middle Eocene Climatic Optimum (MECO), a global warming event that occurred about 40 mya^47^. The subsequent Eocene-Oligocene transition, underscored by Oi-1 glaciation, resulted in a global cooling event^39^ that brought about sea level lowstands^39,41^ and facilitated dispersal of these warm-adapted MRCAs to adjacent refuges via intervening climatically harsh areas. Adaptation to warmer climatic conditions during the MECO thus might have been a major evolutionary advancement in common ancestors showing high dispersal capabilities during early stages of their diversification. Elevated rates of climatic niche evolution in species having broad distributional ranges further support this finding (Supplementary Table 4).

**Figure 4.**
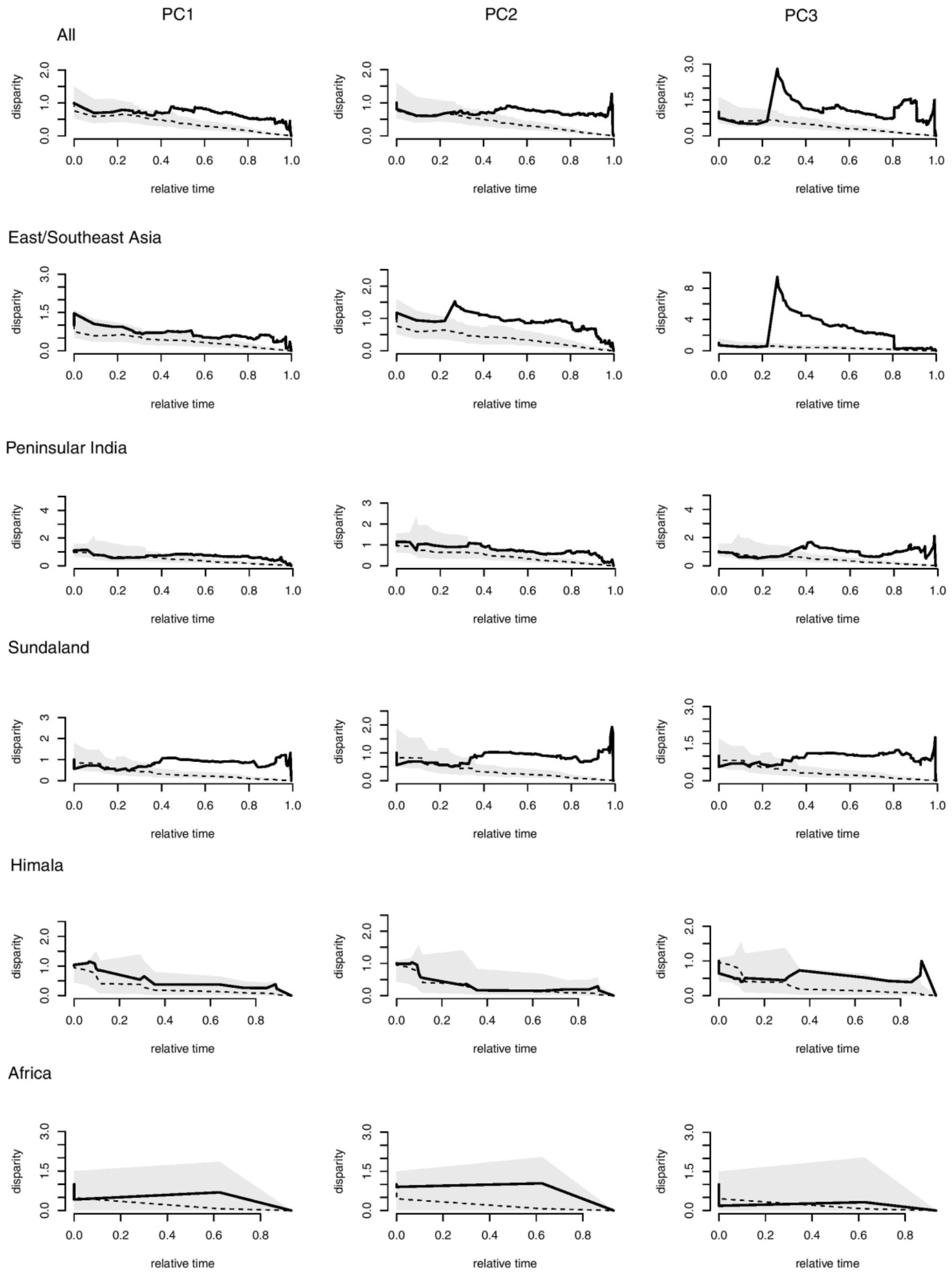
Relative disparity-through-time (DTT) of PC scores representing rhacophorid climatic niches. Solid lines indicate observed DTT values; dashed lines and the corresponding polygons represent the averages and 95% confidence intervals, respectively, of the expectations given a constant accumulation of disparity over time based on 999 pseudoreplicates. Disparity of climatic niche evolution is higher especially during the second half of the diversification in all biogeographic regions except Africa. Markedly higher levels of disparity are observed in PC3 (higher summer temperature) early in the evolution of species from East/Southeast Asia and peninsular India and recently in species from Sundaland.

Variation in rates of climatic niche evolution among biogeographic regions suggests how species have evolved their climatic niches depending on their relative location (Supplementary Table 4). For example, after the Eocene-Oligocene transition, climatic conditions in East/Southeast Asia became colder with the Tibetan-Himalayan orogeny (ca. 30 mya), which, in turn, initiated monsoon cycles^43,44^. Species that continued to occupy cool-wet climates in East/Southeast Asia further adapted towards cooler climatic niches, punctuated by seasonal variation (Fig. 5). In contrast, species that dispersed towards lower latitudes, in particular, Sundaland and peninsular India, invariably had to adapt to climatic niches associated with higher temperatures to survive in warmer tropical environments.

**Figure 5.**
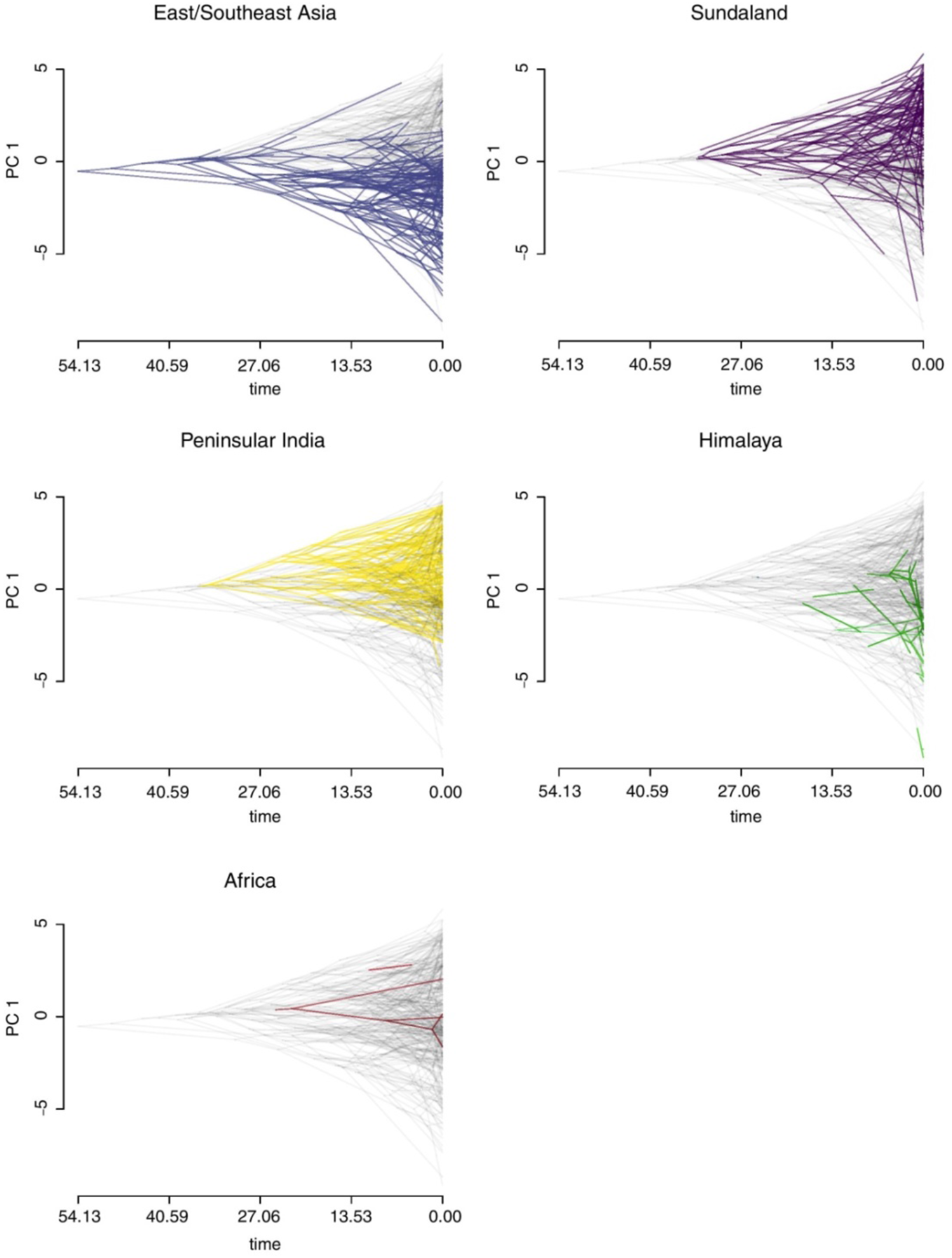
Variation among biogeographic regions in the rate of evolution of the climatic niche explained by the first principal component axis (PC1) of average climatic conditions for each rhacophorid species. The X-axis corresponds to the approximate age of origin of rhacophorid clades to the present (indicated by quartiles); nodes indicate the inferred climatic niches for the most recent common ancestor of the extant taxa defined by that node; positive values along Y-axis represent warmer climates whereas negative values represent colder climates; gray branches in the background of each plot indicate the overall climatic niche evolution of rhacophorids along PC 1. Divergent branches are more frequent near the present time. While species in East/Southeast Asia (blue) and Himalaya (green) have evolved towards colder climatic conditions and those in Sundaland (purple) and peninsular India (yellow) have evolved towards warmer climatic conditions, species in Africa (red) have not deviated much from their ancestral climatic niche.

Island formation during the Miocene^48^ also played a significant role in shaping rhacophorid diversification. In general, a continental climate has characteristics associated with areas within a continental interior, unlike islands, which typically are influenced by surrounding bodies of water, and these differences may affect species composition^49^. Accumulation of a wealth of young rhacophorid lineages in island regions during the Miocene suggests the potential role of islands as species pumps for relatively young species (high SR) as well as refuges (high PD)^50,51^ (Fig. 2b: Q4). In the PCA plot of climatic niche space, island species tend to be associated with intermediate (in subtropical islands) to warmer temperatures (in tropical islands), while mainland species tend to be associated with colder climates influenced by seasonality (Fig. 3). Islands tend to have less-seasonal climates relative to mainland environments^52^, which may buffer island species against the effects of a changing climate.

Climatic stability on islands, however, need not imply optimality. Species dispersing to islands may need to adapt to novel climatic niches. The higher rates of climatic niche evolution for species living on both islands and mainland, compared to only island and only mainland species, suggest that climatic niches evolved extensively in species that show long-distance dispersal (Supplementary Table 5). Interestingly, island species are derived from recent speciation events and have a mainland origin (Supplementary Fig. 3). Island species have higher rates of evolution along PC1 and PC2, whereas mainland species tend to have higher rates along PC3 (Supplementary Table 5). In the context of Rhacophoridae as a whole, the pioneering ancestors of island clades may have been warm-adapted, low-elevation forms that subsequently colonized relatively warm areas (i.e., islands), whereas mainland species that remained in ancestral niches underwent further diversification *in situ*. However, species that successfully colonized islands show higher rates of climatic niche evolution than mainland species (Supplementary Table 5), which suggests that novel ecological opportunities provided by mountains and other complex topographic features within islands may facilitate their high diversity^5,38,53^. According to the theory of niche conservatism, because islands are geographically and environmentally heterogeneous, species may be able to maintain their optimal environment and enhance their chance of persistence simply by dispersing across relatively small distances, especially on topographically complex islands^54^. This may explain how rhacophorids have diversified under less conducive, relatively warmer and less seasonal climatic conditions on islands, by either NE, NC or both. Some of these species, however, might at some point recolonize and re-establish in their continental ancestral area if suitable environmental and ecological conditions return^38,55^ (Supplementary Fig. 3).

Comparison of diversification rates associated with different reproductive modes and among regions (Fig. 6a) reveals that fully aquatic breeders (AQ) are found only in cool-wet East/Southeast Asia^19^, whereas lineages that dispersed from this region to other regions tend to have more terrestrial reproductive modes (direct development, DD; and foam nesting, FN). Moreover, these terrestrial modes are associated with higher diversification rates in all regions. We suggest that a shift from a fully aquatic reproductive mode to more terrestrial modes, but especially DD and FN, facilitated the dispersal of rhacophorids from their centre of origin into optimal but distant climatic oases such as islands. This scenario is corroborated by the pattern of climatic niche occupation of the four reproductive modes: fully aquatic (AQ) and semi-aquatic (GN) species are confined to relatively cool-wet East/Southeast Asia and adjacent subtropical and temperate islands, whereas the more terrestrial modes are confined to warmer climates mostly found in tropical islands (mainly Borneo, Java, Sumatra, Philippines and Sri Lanka), which lie within the intertropical convergence zone (Fig. 6b).

**Figure 6.**
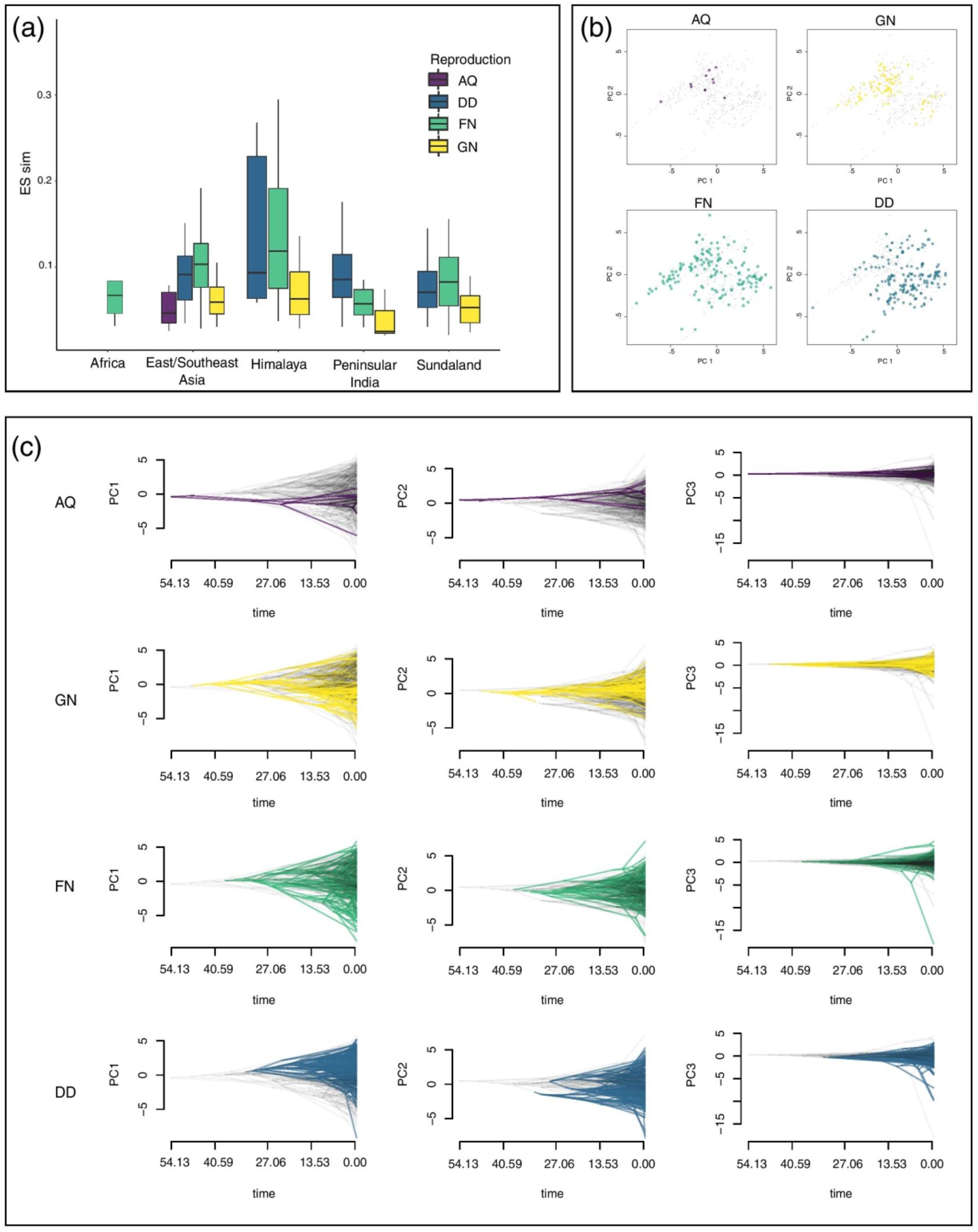
Climatic correlates of different rhacophorid reproductive modes. (a) Box plot showing diversification rates (DR) associated with different reproductive modes in different biogeographic regions (aquatic breeding, AQ; gel nesting, GN; foam nesting, FN; and direct development, DD). Species having FN and DD show high diversification rates. (b) PCA plots depict climatic niche occupation of species that utilize the above four reproductive modes. Aquatic breeders display a conservative (narrow) occupation of climatic niche space, whereas FN have the broadest niche. DD are restricted mostly to warmer but humid and less seasonal climates. (c) Variation in the rate of evolution of the rhacophorid climatic niche explained by the first three principal component axes (PC1–PC3) of average climatic conditions for each reproductive mode. Early AQ shows a pattern of niche conservatism whereas GN, FN and DD, which have facilitated the evolutionary transition of rhacophorids to a more terrestrial life, appear to have evolved their climatic niches during relatively warmer early Eocene (40 mya) conditions. Newly evolved reproductive modes show a pattern of broadening their climatic niches subsequent to the Eocene-Oligocene transition (23–33 mya), which was marked by favourable climatic conditions.

After they colonized East/Southeast Asia, early emerging aquatic-breeding rhacophorids evolved new breeding strategies gel nesting, foam nesting and direct development, which are key evolutionary innovations (KEI) in the clade^18,19^. Species bearing more terrestrial reproductive modes tend to have broader climatic niches, and hence, wider climatic tolerance ranges, which is suggestive of greater dispersal capabilities. For example, rates of climatic niche evolution are higher in FN and DD, whereas the climatic niche in AQ has been more conservative (Supplementary Table 6). FN enables rhacophorids to lay large numbers of eggs in more open and drier habitats, where resistance to desiccation is essential^19^. Indeed, foam-nesters are characterized by large geographic ranges across potentially xeric and open habitats, making them excellent dispersers^19^. Terrestrial DD instead allows frogs to lay eggs in a diversity of humid habitats away from bodies of water. It represents a key innovation that facilitated the evolution of nearly half of all known rhacophorid species^19^, which occupy warmer but humid climatic conditions.

Traitgrams of different reproductive modes suggest that GN may have evolved in association with the MECO global warming event during the early Eocene more than 40 mya^47^ (Fig. 6c). GN, which is characterized by gel-covered, terrestrial eggs and free-living aquatic tadpoles^23^, can be regarded as an initial stage in the evolution of a fully terrestrial life history^19^. Adoption of this reproductive mode has enabled NC-dominant dispersal of rhacophorids into relatively cool-wet areas within close proximity to ancestral climatic niches of AQ (Supplementary Fig. 4).

Traitgrams of the more terrestrial modes FN and DD suggest that they evolved after GN but during the same geological period. Thus, increasing temperatures during MECO may have promoted the adoption of novel modes of reproduction, which equipped rhacophorids for a more terrestrial life. The subsequent Eocene-Oligocene transition, marked by depressed sea levels, land bridge emergence^39,41^ and cooler lowland dry zones, then enabled long-distance dispersal of terrestrial-breeding (FN and DD) lineages. They were successful dispersers insofar as they were able to overcome the ancestral dependence on aquatic habitats while evolving climatic niches (NE) adapted to broader climatic conditions.

The spatial distribution of rhacophorid species is patchy, with islands and some mainland regions having higher diversity and unique assemblages of taxa, which contain older as well as relatively young lineages. This patchiness can be explained in the context of climate, ecological opportunity and key evolutionary innovations. Regions of cool-wet climate may have acted as climatic refuges (high PD regions). They are separated from one another by warmer, climatically harsh low-lying regions and/or sea passages. Lineages overcame these barriers in two main ways: by adapting to harsh climates (NE) or dispersing only during favourable periods (NC). KEIs, such as shifting from a fully aquatic reproductive mode to more terrestrial modes, further facilitated the diversification and dispersal of rhacophorids through these climatically less favourable areas. Finally, ecological opportunity in the form of empty niches seem to have elevated diversification rates on islands, offsetting constraints caused by their less favourable (warmer) climatic conditions. Hence, for rhacophorid frogs, climatic refuges, ecological opportunity, key innovations, long periods of time to adapt to climatic conditions, and climatic niche evolution have combined to promote and sustain a remarkable diversification.

## Methods

Comprising 6% of global amphibian diversity, rhacophorid tree frogs occupy a climatically variable geographic area ranging from tropical and sub-tropical Asia, its continental islands and archipelagos, to Africa. We carried out a series of analyses based on a complete species phylogeny for Rhacophoridae using a combination of phylogenetic inference and phylogenetic imputation to determine how climate has shaped the diversification and dispersal of this charismatic group.

### Phylogenetic inference

Extensive taxon sampling is important in phylogenetic systematics; it increases accuracy and support of evolutionary relationships^25^. Hence, we derived an updated phylogeny of the family Rhacophoridae with the most complete taxon sampling used to date: 415 extant species representing all 22 genera. Although previous studies have resolved many of the earlier controversies in rhacophorid phylogeny (Table S1), taxonomic discordances persist^18^, hindering the testing of hypotheses for major evolutionary questions. Moreover, the genetic data available for different rhacophorid lineages are uneven, ranging from none for some species to whole-genome sequences for others, which leads to incomplete taxon sampling. Therefore, we took three main steps to derive a complete species-level phylogeny: (i) establish a phylogenomic backbone, (ii) compile Sanger sequence data of non-chimeric sequences, and (iii) add species without genetic data using phylogenetic imputation^56^. We began by reanalysing the anchored hybrid enrichment dataset (AHE) of Chen et al. (2020)^18^. Although there was strong support for most branches of the rhacophorid tree, one clade (comprised of *Pseudophilautus, Kurixalus* and *Raorchestes*) proved recalcitrant; it was unstable in species-tree analyses. We estimated Maximum Likelihood trees for each locus using RAxML 8.2.12^57^; branch support for each gene tree was based on 50 bootstrap replicates. Preliminary analyses indicated that the source of instability was the position of *Nasutixalus*, given that a species tree without that terminal using ASTRAL III v.5.7.3^58^ yielded 100% local posterior probabilities for the relationship among *Pseudophilautus, Kurixalus* and *Raorchestes*. We therefore enumerated the loci that supported each of the three alternative topologies, as well as the corresponding average bootstrap values of the loci supporting each of them. Given that one of the alternative topologies tended to be more strongly supported by loci with higher average bootstraps (Fig. S5), that topology was chosen for downstream analyses.

Once a stable backbone tree was obtained, we compiled all Sanger sequence data available from GenBank (last accessed January 2021), which included 315 rhacophorid species (number of species/genus in parentheses): *Beddomixalus* (1), *Buergeria* (3), *Chirixalus* (2) *Chiromantis* (3), *Feihyla* (4), *Ghatixalus* (3), *Gracixalus* (9), *Kurixalus* (17), *Leptomantis* (10), *Liuixalus* (6), *Mercurana* (1), *Nasutixalus* (3), *Nyctixalus* (3), *Philautus* (31), *Polypedates* (14), *Pseudophilautus* (56), *Raorchestes* (59), *Rhacophorus* (32), *Rohanixalus* (2), *Taruga* (3), *Theloderma* (25) and *Zhangixalus* (28; see supplementary material, Table S7). Sequences were obtained for seven gene fragments: mitochondrial loci 16S, 12S and *cyt-b*; and nuclear loci *BDNF, Rag-1, rhod* and *Tyr*. Each locus was aligned separately using MUSCLE^59^ as implemented in MEGA v.6^60^. Sequences of all fragments were then concatenated, with a total alignment length of 3923 base pairs (bp).

We obtained complete species-level trees by using the two-stage Bayesian approach PASTIS^56^. This method uses as inputs a backbone topology based on molecular data, a set of taxonomic postulates (e.g., constraining species to belong to specific genera or families) and user-defined priors on branch lengths and topologies. Based on these inputs, PASTIS produces input files for MrBayes 3.2.5^61^, which generates a posterior distribution of complete ultrametric trees that capture uncertainty under a homogeneous birth-death prior model of diversification and placement constraints. We used PASTIS version 0.1-2, with functions from the APE 5.4^62^and CAPER 0.2^63^ packages. There are two main assumptions to this approach: (i) taxonomic groups (e.g., genera) are monophyletic unless there is evidence (i.e., genetic data) that suggests otherwise; and (ii) reasonable edge-length and topology priors (i.e., birth-death models) exist. PASTIS has been used to provide large-scale trees of several higher-level taxa, including birds^64^, squamates^65^, and sigmodontine rodents^66^. Thomas et al. (2013)^56^ categorize species into three types: type 1 species have genetic information (i.e., genomic data + Sanger sequencing data); type 2 species lack genetic information but are congeners of a species with genetic information; and type 3 species lack genetic data and are members of a genus that lacks genetic data. In our dataset we have 321 (genetic data present) and 94 (genetic data absent; Figure S6) species exclusively from types 1 and 2, respectively. Our backbone tree was established using the anchored hybrid enrichment dataset^18^. Species without genetic data are constrained to their closest relatives based on morphology as indicated in published literature (Table S8) and the best partitioning scheme for the Sanger dataset was determined using PartitionFinder2^67^. The general time-reversible model with an invariable gamma rate for each partition, birth-death as prior probability distribution on branch lengths, fixed extinction-rate priors and exponential net speciation-rate priors were assigned to the alignment and constraints was run on MrBayes 3.2.5 for 20 million generations. Convergence was assessed by inspecting the log-output file in TRACER v.1.6^68^ and by ensuring ESS values were greater than 200. The first 10% of the trees were discarded, and the post burn-in trees used to infer the maximum clade credibility tree using TREEANNOTATOR v.1.10.4^69^. The maximum clade credibility tree, as well as a set of 1000 post burn-in topologies (provided in supplementary data), were retained for further analyses (see below).

To examine the temporal context of divergence and correlate it with geological and climatic events, we estimated divergence times among lineages in a separate run of the above partitioned dataset. We initially used the lognormal relaxed clock model with default clock rate priors and, subsequently, the default strict clock model. The lognormal relaxed clock model, which showed the greatest fit, was used for further analysis. We calibrated the tree using two literature-based, molecular-estimated points from Meegaskumbura et al. (2019)^38^ and Chen et al. (2020)^18^ (viz., the age of the MRCA of extant *Pseudophilautus*, 21.93–45.14 mya; and the age of the MRCA of clade A, 31.65–40.53 mya, respectively).

### Testing for variation in rates of lineage diversification

We tested for potential correlates of speciation rates using ES-sim, a semi-parametric test for trait-dependent diversification analyses^70^. Instead of modelling the relationship between traits and diversification, ES-sim tests for correlations between summary statistics of phylogenetic branching patterns and trait variation at the tips of a phylogenetic tree. It uses the DR statistic^65^, a non-model-based estimator of speciation rate that is computed for a given species, as a weighted average of the inverse branch lengths connecting the focal species to the root of the phylogeny (e.g., the root-to-tip set of branches). The use of tip-specific metrics of speciation rate may provide an alternative to parametric state-dependent diversification due to the elevated rates of false-positive results, given that heterogeneity in diversification rates of the underlying phylogeny could bias inferences of associations between traits and diversification regardless of their underlying relationship^71^. Simulations show that the use of ES-sim for continuous traits shows equal or superior power than QuaSSE^70^. In addition, given that they are computationally efficient, tip-specific metrics make it feasible to explore the impact of phylogenetic uncertainty. ES-sim was implemented using the code provided by Harvey & Rabosky (2018)^70^ (available at https://github.com/mgharvey/ES-sim) and we initially assessed variation in speciation rates among lineages by mapping variation in the DR statistic along the maximum clade credibility tree using the ‘contmap’ function in PHYTOOLS 0.7-47^72^. Given that the DR statistic tends to focus more on processes closer to the present, we compare those results with lineages-through-time (LTT) plots using 1000 alternative trees using the ‘mltt.plot’ function in APE.

### Spatio-temporal patterns of rhacophorid diversification

We downloaded distribution ranges as spatial data polygons from IUCN (2020)^36^ http://www.iucnredlist.org/technical-documents/spatial-data for all available rhacophorid species and transformed them into a presence-absence matrix (PAM) by using a 1°×1° global grid using the function ‘lets.presab’ in R package letsR V 3.2^37^. Species richness (SR) was visualized as species richness values per cell on a world map by applying the function ‘plot’ in letsR. Using the maximum clade credibility tree, we calculated phylogenetic diversity (PD), determined as the sum of all co-occurring species branch lengths^73^ within each cell, by applying the function ‘lets.maplizer,’ and obtained a raster to map geographic patterns of PD. Similarly, geographic distribution maps were created for both the DR estimated above^70^ and species crown age (SA), estimated as the length of the terminal branch subtending each species until the most recent speciation event, to visualize spatial patterns of diversification.

As many young (recent) lineages and close relatives attain high values of DR, and those with few close relatives (older/basal lineages) yield lower values (Fig. 1), DR allows us to effectively score all lineages with respect to their relative phylogenetic isolation^3^. As the above metrics (SR, PD, DR and SA), especially DR and SA, are significantly correlated (Fig. S1), the spatial analyses produce more or less similar patterns regardless of the metric assessed. In the main text, we therefore focus on results using DR values. Following Kennedy et al. (2017)^3^, we used the species ranks of DR to divide our distributional database into quartiles. The first quartile contains the oldest and least diverse lineages, while the fourth quartile contains the youngest and most diverse ones (Figure 2B). We subsequently generated maps of species richness at the 1°×1° scale for each quartile using the ‘lets.maplizer’ function in the package letsR. This enables us to visualize areas that have accumulated a higher or lower number of species in each DR quartile. Alternatively, this reveals geographic variation in lineage accumulation through space and time. Moreover, relative and distinct increase in diversifications within the same regions in quartiles would support the idea of NC-dominant early dispersal and diversification. Finally, to explicitly test whether islands serve as species pumps, we used island/mainland as a binary trait and ran ES-sim on 100 potential topologies. Due to the highly correlated nature of DR and SA (Fig. S7), it can be assumed that each quartile represents the temporal aspect of diversification as well.

To further investigate species accumulation through time in different biogeographic regions (East/Southeast Asia, Sundaland, peninsular India, Himalaya and Africa), we used subsets of the rhacophorid phylogeny based on geographic region and assigned birth-death models of diversification under four conditions for the birth and death rates specified in the package RPANDA^74^. Best models were selected for each biogeographic region based on AICc values, and diversity through time was visualized using the function ‘plotdtt’ in RPANDA (Figure S2).

### Ecological and biogeographic data

Information on geographical distribution of rhacophorid species derived from occurrence records was obtained from the GBIF database (https://www.gbif.org) using RGBIF 1.4.0^75^ in R v.3.6.1^76^. Distribution data for species not represented in GBIF were obtained directly from the literature and IUCN species range maps. The final distribution dataset comprised 3846 geographical coordinates of all species. Nearly 65% of those species (N = 273) were represented by single-occurrence records. This is not uncommon in compilations of this nature^9,33^ and reflects the high degree of local endemism of rhacophorid species. Information on 19 bioclimatic variables for each occurrence point were obtained from WORLDCLIM 2.0^32^ using the “extract” function in RASTER 3.0-7^77^. Mean values for each bioclimatic variable for each species are provided in Table S4. We then used the average bioclimatic conditions in a principal components analysis (PCA) based on their correlation matrix. The axes to be retained for further analyses were determined using the broken-stick method as implemented in VEGAN 2.5-6^78^. We assume that the measured species means are a reasonable approximation of the realized climatic niche of the species^9,33^.

### Climatic correlates of rhacophorid diversification

When implementing ES-sim^70^, we used 100 simulations to build the null distribution of trait-speciation associations for significance testing. The designated potential correlates of speciation rates were the first three principal components of the climatic niches, mean elevation and whether these species were island endemics. To account for phylogenetic uncertainty, we repeated each analysis for 100 potential topologies. Analyses were carried out separately for each potential correlate (Table S3).

We assessed the extent to which species traits had accumulated over time in each biogeographic region (i.e., climatic niche evolution; NE along each PC axis) by using disparity-through-time (DTT) plots^79^, with expected disparities calculated based on 1000 resamplings using the ‘dtt’ function in GEIGER 2.0.6.2^80^ and with phenograms (projections of the phylogenetic tree in a space defined by phenotype and time) using the ‘phenogram’ function in PHYTOOLS. Biogeographic categorizations for these analyses are based on Chen et al. (2020)^18^. Here, East/Southeast Asia includes mainland China and subtropical islands such as Hainan, Taiwan and Japan; Sundaland includes peninsular Malaysia and islands such as Borneo, Sumatra, Java and Philippines; and peninsular India includes mainland India and Sri Lanka.

We tested whether evolutionary rates of the main climatic niche axes (PC1, PC2 and PC3) differ significantly among species inhabiting different biogeographic regions, islands, mainland, and both islands and mainland. We assumed that species living both on islands and mainland have a broader climatic niche than only-island and only-mainland species; high rates of climatic niche evolution of those species would suggest NE. We used 1000 potential trait histories from stochastic character mapping and then fit two alternative models of evolution on each studied trait—one that fixes the rate of niche evolution to be identical between island / mainland inhabitants, and an alternative model in which the inhabitants have separate rates. We calculated the Akaike Information Criterion for small sample size (AICc) from the maximum likelihood estimate on each tree using the ‘brownie.lite’ function in PHYTOOLS. Subsequently, we calculated model-averaged estimates of evolutionary rates for each category. Similarly, rates of climatic niche evolution associated with different reproductive modes were assessed using the above method.

Finally, to infer the biogeographic history of Rhacophoridae by determining whether lineages originated on islands or on mainland, a model-testing approach was applied using the R package BioGeoBEARS v.0.2.1^81^ based on the maximum credibility tree. Species occurrences were categorized according to the biogeographic areas modified from Chen et al., 2020^18^: (1) Africa mainland; (2) peninsular India (mainland); (3) Sri Lanka (island); (4) East/Southeast Asia mainland; (5) East/Southeast Asian islands, including Japan, Taiwan, Hongkong and Hainan; (6) Sundaland (island archipelago); and (7) Himalaya. Given the extreme geological complexity of the region through time, dispersal constraints were not applied^18^. Analyses used six biogeographic models specified in the package (Table S9) and model fit was assessed using the Akaike Information Criterion (AIC) and Akaike weights^81^.

